## Supplementary material for "Contrasting responses to aridity by different-sized decomposers cause similar decomposition rates across a precipitation gradient": Table S1, Table S2

**Supplementary information**

**Table S1** – Results of post hoc pairwise comparisons between macro-decomposer assemblages across experimental sites (AV – Avdat, BG – Bet Guvrin, HS – Havat Shikmim, MS – Meishar, NS – Nahal Shita, SS – Sayeret Shaked, RH – Ramat Hanadiv). P-values in the right column are adjusted according to the Benjamini-Hochberg procedure.

| **Comparison** | | | **Df** | **Sum of squares** | **F** | **R^2^** | **P-value** | **Adjusted P-value** |
| --- | --- | --- | --- | --- | --- | --- | --- | --- |
| SS | vs | AV | 1 | 3.05 | 8.64 | 0.10 | 0.001 | 0.00105 |
| SS | vs | HS | 1 | 2.75 | 7.58 | 0.09 | 0.001 | 0.00105 |
| SS | vs | BG | 1 | 3.37 | 9.61 | 0.12 | 0.001 | 0.00105 |
| SS | vs | RH | 1 | 6.67 | 21.74 | 0.24 | 0.001 | 0.00105 |
| SS | vs | MS | 1 | 3.26 | 8.99 | 0.11 | 0.001 | 0.00105 |
| SS | vs | NS | 1 | 3.41 | 9.31 | 0.12 | 0.001 | 0.00105 |
| AV | vs | HS | 1 | 2.85 | 7.19 | 0.09 | 0.001 | 0.00105 |
| AV | vs | BG | 1 | 3.42 | 8.91 | 0.12 | 0.001 | 0.00105 |
| AV | vs | RH | 1 | 5.63 | 16.64 | 0.20 | 0.001 | 0.00105 |
| AV | vs | MS | 1 | 1.47 | 3.72 | 0.05 | 0.001 | 0.00105 |
| AV | vs | NS | 1 | 1.64 | 4.08 | 0.06 | 0.001 | 0.00105 |
| HS | vs | BG | 1 | 2.49 | 6.30 | 0.09 | 0.001 | 0.00105 |
| HS | vs | RH | 1 | 4.93 | 14.07 | 0.18 | 0.001 | 0.00105 |
| HS | vs | MS | 1 | 2.39 | 5.88 | 0.08 | 0.001 | 0.00105 |
| HS | vs | NS | 1 | 2.34 | 5.66 | 0.08 | 0.001 | 0.00105 |
| BG | vs | RH | 1 | 4.81 | 14.35 | 0.19 | 0.001 | 0.00105 |
| BG | vs | MS | 1 | 3.09 | 7.84 | 0.10 | 0.001 | 0.00105 |
| BG | vs | NS | 1 | 2.75 | 6.86 | 0.10 | 0.001 | 0.00105 |
| RH | vs | MS | 1 | 5.39 | 15.43 | 0.19 | 0.001 | 0.00105 |
| RH | vs | NS | 1 | 4.89 | 13.82 | 0.18 | 0.001 | 0.00105 |
| MS | vs | NS | 1 | 0.97 | 2.36 | 0.03 | 0.002 | 0.002 |

**Table S2** – Dissimilarity matrix between macro-decomposer assemblages of the different site-season combinations. Cells are filled on a green-yellow-orange-red scale with increasing Bray-Curtis dissimilarity.

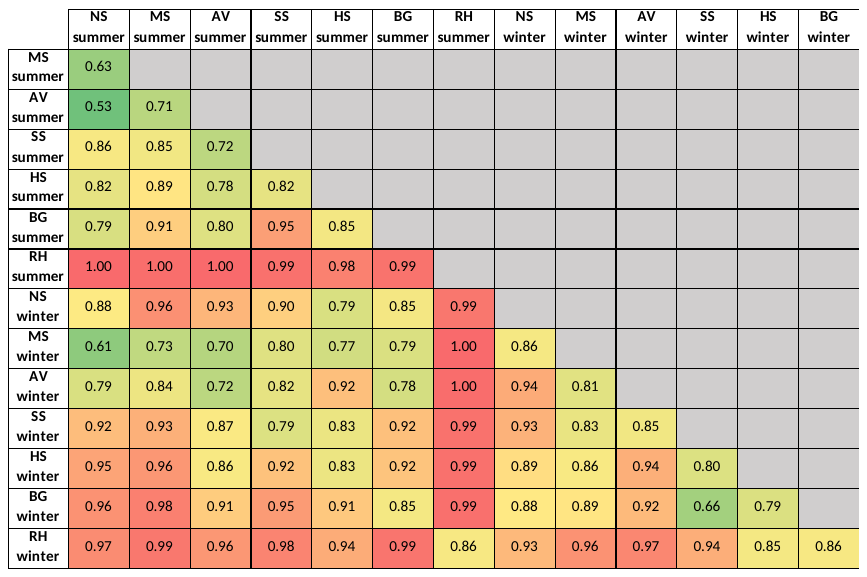
